# Rapid and Quantitative Phage Susceptibility Test by Ramanome

**DOI:** 10.64898/2026.02.07.704537

**Authors:** Xiao Han, Xiaofu Wan, Yang Zhou, Xiaoting Fu, Xiaoshan Zheng, Bo Gao, Shi Huang, Anle Ge, Jiadong Huang, Hongzhou Lu, Jian Xu

## Abstract

Antimicrobial resistance poses an escalating global threat, renewing interest in bacteriophage therapy as a precision alternative to antibiotics. However, clinical translation remains hindered by the lack of rapid and quantitative phage susceptibility testing (PST) platforms capable of evaluating host range, infection potency, and effective multiplicity of infection (MOI). Here we present RPST, a ramanome-based phenotypic platform that captures infection-induced remodeling of bacterial macromolecular composition to unify these diagnostic requirements within a single workflow. RPST integrates four Raman biomarkers into a Composite Infection Index (CII), enabling rapid and lysis-independent discrimination between susceptible and resistant bacterial populations within ∼1 hour, with 96.0% categorical concordance (24/25) to plaque assays. As a continuous population-level metric, CII quantifies the proportion of infected cells, allowing quantitative ranking of phage potency against shared hosts. By resolving CII trajectories across the MOI and time, RPST further determines the minimal effective MOI, which is the lowest phage-to-bacterium ratio sustaining self-propagating infection, thereby defining the lower boundary for therapeutic feasibility. Together, these capabilities transform PST from static compatibility assays into a dynamic and quantitative framework that bridges *in vitro* infectivity assessment and infection dynamics relevant to phage therapy.

**Impact Statement:** Based on the rapid emergence of antimicrobial resistance, this study introduces RPST, a novel ramanome-based phage susceptibility testing platform. RPST detects phage-induced biochemical remodeling in bacteria within ∼1 hour, achieving 96.0% concordance with gold-standard plaque assays. By integrating four Raman biomarkers into a Composite Infection Index, it not only distinguishes susceptible from resistant strains but also quantifies phage potency and determines the minimal effective multiplicity of infection required for self-sustaining infection. This transformative approach moves beyond binary diagnostics to offer a dynamic, quantitative framework for precision phage therapy, significantly accelerating therapeutic decision-making and enhancing our ability to combat resistant infections.

## Introduction

Antimicrobial resistance (AMR) remains a critical global health threat. In 2019, it caused over one million deaths, and the situation is worsening as effective antibiotic options continue to diminish (1, 2). The World Health Organization projects that by 2050, AMR could lead to over 10 million deaths annually, surpassing cancer as the leading cause of mortality (1). In this context, phage therapy has re-emerged as a promising countermeasure. Bacteriophages, the natural viruses that target bacteria, have unique therapeutic advantages, including high specificity, self-amplifying capacity, and minimal disruption to the host microbiome (3, 4). They directly eradicate target pathogens through lytic cycles and have shown remarkable efficacy against multidrug-resistant bacterial strains and biofilm-associated infections where antibiotics often fail (5). This has spurred a surge of interest in phage therapy, with over 40 clinical trials currently registered on ClinicalTrials.gov that target a wide range of infections (6, 7).

Despite its promise, clinical translation of phage therapy faces significant challenges, leading to inconsistent therapeutic outcomes (8). This variability is largely due to three major obstacles: the intrinsically narrow host range of individual phages (8), the inevitable emergence of bacterial resistance driven by the co-evolutionary dynamics between phages and their hosts (3), and the difficulty of achieving therapeutically relevant phage concentrations at infection sites, where effective clearance requires phage replication to outpace bacterial proliferation (9). Addressing these barriers critically depends on the development of robust and clinically informative phage susceptibility testing (PST) platforms (10). An ideal PST system must therefore (*i*) rapidly and accurately determine whether a phage can infect a given pathogen, thereby resolving the constraint of narrow host range; (*ii*) discriminate the relative potency of active candidates, enabling the selection of phages most capable of countering resistance; and (*iii*) evaluate whether infection dynamics, including phage replication and secondary infection, can sustain effective bacterial clearance under clinically relevant multiplicity of infection (MOI) conditions, ensuring therapeutic feasibility (11).

Current PST strategies can be broadly categorized as genotypic or phenotypic, yet neither satisfies the three essential diagnostic criteria. Genotypic methods, which quantify phage genome copies, offer high sensitivity and rapid single-endpoint readouts. Although they can indirectly estimate infection potential, they do not measure actual infectivity, are prone to false positives and negatives, and lack the temporal resolution needed to capture infection dynamics, limiting their clinical relevance (12). Among phenotypic approaches, growth-based assays such as plaque assays (the current gold standard), disk diffusion, broth microdilution, and optical density (OD) monitoring can reliably distinguish susceptible from resistant hosts, but are time-consuming, offer only qualitative assessments of infection potency, and fail to resolve dynamic infection processes (8, 10, 13). Metabolism-based assays, including isothermal microcalorimetry and probe-coupled flow cytometry, can shorten turnaround time and improve throughput, yet typically yield static, qualitative outputs that fail to simultaneously capture infectivity, relative potency, and phage replication dynamics (8, 11). Consequently, the development of a rapid, accurate, quantitative, and dynamic PST system remains a critical unmet need. To contextualize these unmet needs, representative PST strategies and their diagnostic capabilities are compared (**Table 1**), which suggests that no existing methods simultaneously resolve infectivity, relative lytic potency, and infection sustainability under defined MOI conditions.

**Table 1.** Comparison of representative phage susceptibility testing (PST) strategies and their diagnostic capabilities.

| Method / Platform | Principle | Key advantages | Major limitations | Ref |
| --- | --- | --- | --- | --- |
| Plaque assay | Plaque formation | High accuracy; direct evidence of infectivity; providing plaque morphology | Time-consuming; qualitative or semi-quantitative; endpoint-only | (10) |
| qPCR | Phage genome copy number | Rapid; highly sensitive; quantitative | Unable to measure infectivity; unable to distinguish productive from abortive infection; prone to false positives/negatives | (12) |
| Optical density (OD) monitoring / kill curves | Bulk growth inhibition | Dynamic monitoring; relatively rapid | Unable to distinguish live/dead cells; confounded by aggregation and growth arrest; indirect and non-quantitative infectivity | (8, 11) |
| Disk diffusion | Growth inhibition zones | Simple; standardized | Time-consuming; not quantitative; lacking kinetic information | (13) |
| Flow cytometry-based | Membrane integrity | Rapid; high-throughput; single-cell resolution | Non-specific signals (false positives); complex assay design; unable to distinguish productive from abortive infection | (47) |
| Isothermal microcalorimetry | Heat flow from metabolism | Broadly applicable; dynamic | Time-consuming; indirect metabolic readout via thermal signals | (7) |
| RPST (this work) | Population-level biochemical remodeling | Rapid; quantitative; simultaneously resolving infectivity, lytic potency, and minimal effective MOI | Currently population-averaged; requiring multi-center validation | - |

Ramanomics has emerged as a new analytical platform in phage research (14). For example, surface-enhanced Raman spectroscopy (SERS)-based probes have enabled the specific identification and discrimination of pathogenic microorganisms (15–18), while spectral profiling of infected cells and biofilms has advanced the understanding of phage-bacterium interactions (19–21). The integration of tip-enhanced Raman spectroscopy (TERS) further achieves nanoscale spatial resolution, allowing visualization of viral surface topology and bacterial adhesion proteins (22). Owing to its rapid, label-free, and non-destructive molecular fingerprinting capability (23), which captures a rich spectrum of cellular metabolic phenomes (24, 25), ramanomics holds strong potential for probing infection-induced biochemical remodeling.

Here, a *ramanome* (26) refers to a structured collection of spontaneous Raman spectra acquired from a certain number of cells within an isogenic population under a defined condition and time window, collectively representing a population-level metabolic phenome. Each individual Raman spectrum encodes thousands of vibrational features associated with cellular macromolecules, such that the ensemble captures the composite biochemical state of the population rather than single-molecule signatures. Importantly, intrinsic cell-to-cell heterogeneity is preserved within a ramanome and contributes biologically meaningful variation rather than experimental noise. In the context of phage infection, ramanomes primarily monitor host bacterial cells, capturing coordinated metabolic remodeling that emerges during the latent phase prior to overt lysis.

Indeed, recent studies have shown that spontaneous Raman spectroscopy can distinguish infected from uninfected bacterial cells by detecting alterations in major macromolecular components within the fingerprint region (27, 28). However, these studies have largely focused on isolated phage-host pairs, yielding case-specific spectral changes rather than broadly applicable diagnostic principles. This fragmentation reflects the extensive genetic and physiological diversity among phages (29), which drives heterogeneous host responses and hampers the establishment of a unified Raman-based susceptibility testing framework. To overcome these limitations, a systematic and expandable ramanome-based framework is needed to enable rapid, quantitative, and dynamic assessment of phage susceptibility across diverse phage-host systems.

Here we developed a ramanome-based phage susceptibility test (RPST) that harnesses infection-induced remodeling of bacterial macromolecular composition as a rapid phenotypic readout of phage-host interactions. By capturing coordinated changes in nucleic acids, proteins, and lipids at the population level, RPST enables rapid and lysis-independent discrimination between phage-susceptible and phage-resistant bacterial populations. Beyond binary discrimination, RPST provides a quantitative framework to evaluate infection efficacy across different MOI and to resolve differences in lytic potency among phages against the same host. In addition, by integrating single-endpoint analysis with time-resolved ramanome trajectories, RPST captures infection progression and the capacity of phage populations to sustain propagation over time. Together, these features position RPST as a rapid, quantitative, and dynamic phenotypic platform that addresses the core diagnostic requirements of phage susceptibility testing, bridging static compatibility assessment and infection dynamics relevant to therapeutic feasibility.

## Results

### Overview of the RPST Workflow

The conventional plaque assay, the clinical gold standard for phage susceptibility testing (8), requires multiple labor-intensive steps, including isolate expansion, phage-host co-incubation, double-layer plating, incubation, and plaque enumeration, typically taking 24∼36 hours to determine host range (**Fig. 1A**). In contrast, RPST leverages phage-induced remodeling of bacterial macromolecular composition, captured by Raman spectroscopy, to classify infection outcomes within approx. 1 hour (**Fig. 1B**).

**Figure 1.**
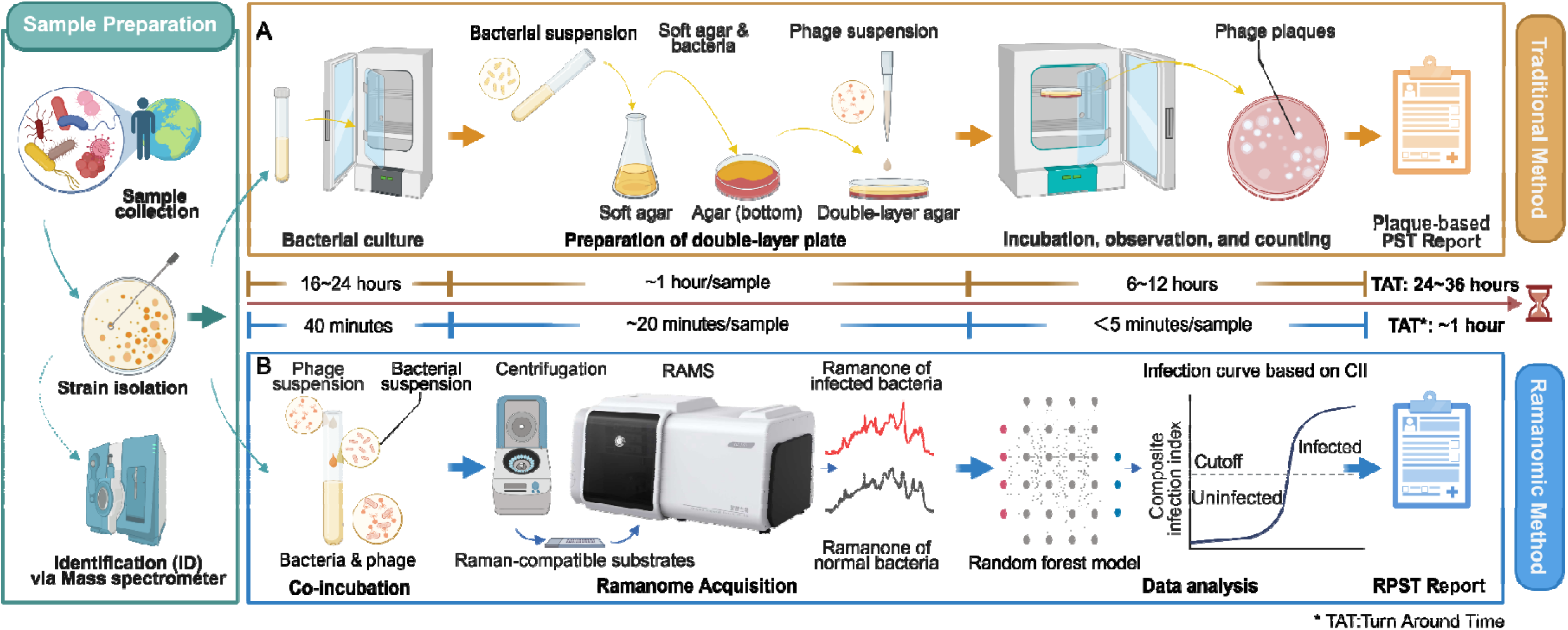
Overview of Ramanome-Based Phage Susceptibility Testing (RPST). (**A**) Conventional workflow (upper panel). Bacterial colonies isolated from environmental or clinical samples are enriched on solid agar plates (12∼16 hours). Susceptibility testing via plaque assays (double-layer agar method) involves three steps: (*i*) Bacterial colonies are isolated and enriched on solid agar plates for 12∼16 hours; (*ii*) Bacterial suspensions are co-incubated with various phages, followed by a two-layer plate assay; (*iii*) Plaque observation after 6∼12 hours of incubation. Total detection time: 24∼36 hours. (B) RPST workflow (lower panel). (*i*) Bacterial colonies are suspended and co-incubated with phages for 40 minutes; (*ii*) Centrifugal washing and transfer to Raman-compatible substrates; (*iii*) Acquisition of phage-bacteria interaction spectra (≥ 60 spectra/sample) followed by standardized analysis. Total detection time: ∼1 hour. Created with BioRender.com.

The RPST workflow consists of three sequential steps: (*i*) sample preparation, involving co-incubation of bacterial suspensions with phages for 40 minutes; (*ii*) ramanome acquisition, including washing, deposition onto Raman-compatible substrates, and spectral collection; and (*iii*) data analysis, in which a standardized processing pipeline computes a composite infection index (CII) for susceptibility determination (**Fig. 1B**).

### Identification of Phage-Induced Biomarkers by Raman Fingerprints to Distinguish Infected from Uninfected Bacterial Cells

Previous studies have demonstrated that Raman spectroscopy can capture infection-induced alterations in cellular biomolecules (27, 28), However, the biochemical diversity of phage-host interactions complicates the identification of conserved diagnostic features. To establish a robust ramanome-based framework for PST, we sought to identify spectral biomarkers that consistently distinguish infected from uninfected bacterial populations across representative systems.

To this end, we analyzed three representative phage-host systems (*E. coli* ATCC11303-T1, ATCC11303-T4, and ATCC25922-T4) chosen to sample both phage- and host-dependent variation (T1 is FhuA-dependent and relatively host-restricted, whereas T4 is a broadly lytic *Myoviridae* phage (30)). Ramanomes were profiled at 20∼120 min (20-min intervals; **Fig. S1A-C**) to capture the trajectory from early adsorption and macromolecular remodeling to population-level lysis, ensuring candidate markers reflect sustained infection dynamics rather than transient noise. For each fingerprint peak, the area under the receiver operating characteristic curve (AUC) was computed between untreated and infected groups (MOI = 10). Peaks with mean AUC values > 0.80 at four or more time points across all three susceptible systems were retained, yielding 15 candidate spectral regions (86 wavenumbers; **Fig. 2A, top**). Differential spectral analysis further refined these candidates to four conserved regions exhibiting consistent directional changes: 658∼663 cm¹ (C–S stretching in cystine, ↓), 713∼719 cm¹ (pyrimidine ring breathing in nucleic acids, ↓), 1430∼1450 cm¹ (CHC bending in lipids and protein, ↑), and 1589∼1595 cm¹ (C=C skeletal vibration in aromatic amino acids, ↑; **Fig. 2A**, **bottom**; **Supplementary Table1**). These conserved signatures form the basis of subsequent RPST metric construction.

**Figure 2.**
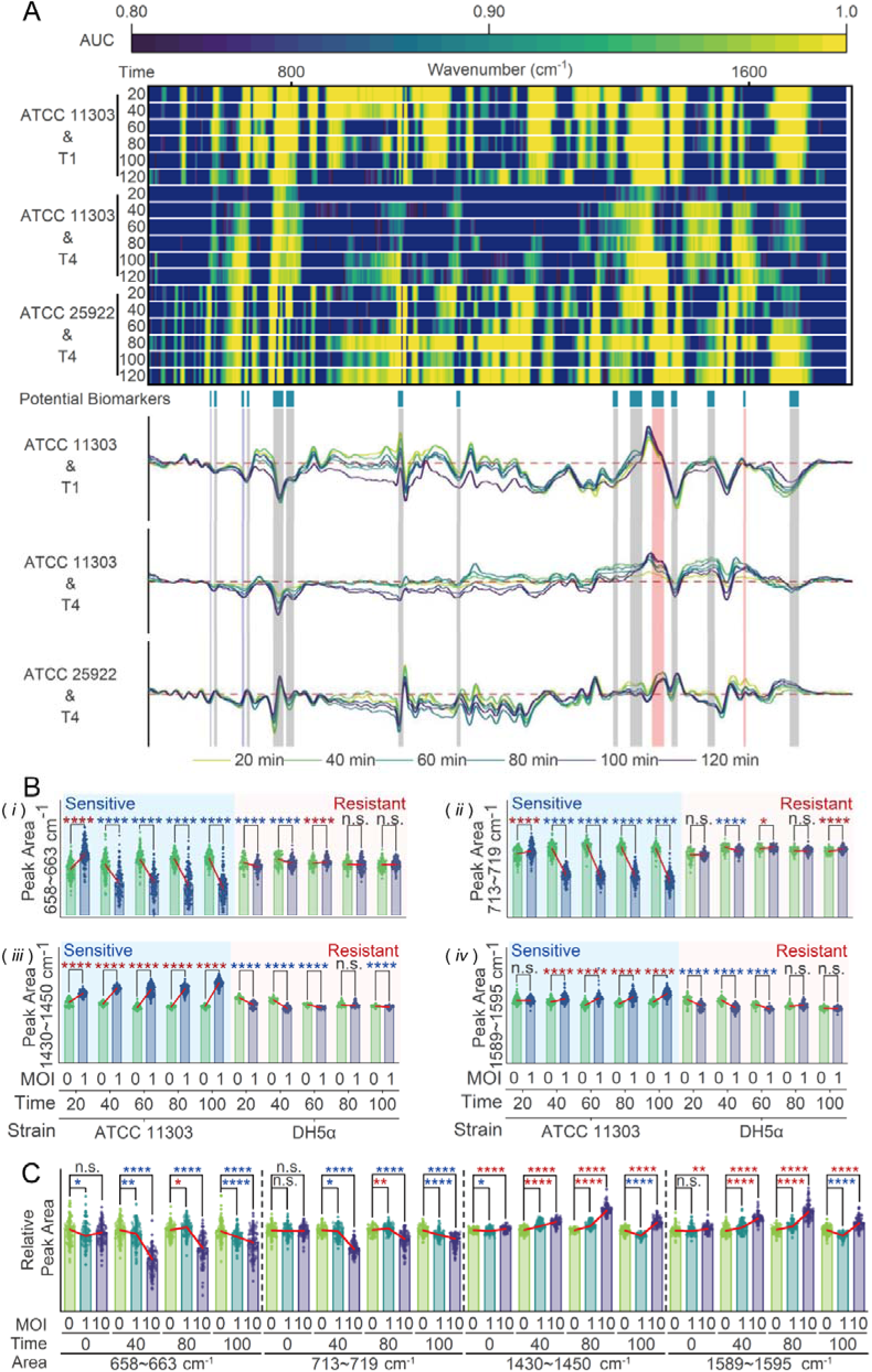
Identification of infection-associated Raman biomarkers. (**A**) ROC analysis was applied to Raman spectral fingerprint regions to identify biomarkers distinguishing infected from uninfected bacteria. Raman spectra were collected from three sensitive systems (T1-*Escherichia coli* ATCC11303, T4-*E. coli* ATCC11303, T4-*E. coli* ATCC25922) at six time points (20∼120 min) under MOI = 10. Peaks showing an average area under the ROC curve (AUC) > 0.80 in at least four of six time points across all three systems are considered candidate biomarkers (n = 15). Combined with difference spectrum analysis, only those showing consistent directional changes (either increased or decreased upon infection) across systems are retained, resulting in four universal candidate regions: 658∼663 cm¹ (decreased), 713∼719 cm¹ (decreased), 1430∼1450 cm¹ (increased), and 1589∼1595 cm¹ (increased). (**B**) Temporal dynamics of the four biomarkers are compared between sensitive and resistant systems. In the sensitive T1-ATCC11303 system, 658∼663 cm¹ and 713∼719 cm¹ gradually decrease, whereas 1430∼1450 cm¹ and 1589∼1595 cm¹ continuously increase over time. In contrast, in the resistant T1-DH5α system, the initial decrease of 658∼663 cm¹ and 713∼719 cm¹ (≤ 40 min) is followed by recovery or stabilization, while 1430∼1450 cm¹ and 1589∼1595 cm¹ transiently decrease at early time points (≤ 60 min) and then remain unchanged or slightly decline. (**C**) Temporal dynamics of the four biomarkers in the weakly sensitive system under different MOI conditions. Under MOI = 10, 658∼663 cm¹ and 713∼719 cm¹ consistently decrease, while 1430∼1450 cm¹ and 1589∼1595 cm¹ increase. Under MOI = 1, responses fluctuate: the first two regions decrease at 40 min, increase at 80 min, and sharply drop at 100 min; the latter two increase at 40∼80 min but also decline at 100 min. n.s., not significant; \**p* < 0.05, \*\**p* < 0.01, \*\*\**p* < 0.001, \*\*\*\**p* < 0.0001.

To evaluate whether these spectral regions can discriminate phage-sensitive from resistant strains, we collected ramanomes from *E. coli* ATCC11303 (T1-sensitive) and DH5α (T1-resistant) after infection with phage T1 (MOI = 1) across 20∼100 min, with uninfected cells (MOI = 0) as controls (**Fig. S1D, F**). In ATCC11303, global spectral divergence from controls becomes evident at 40 min and intensifies over time, whereas DH5α displays only transient deviations (< 60 min) that largely subside by 80 min. Analysis of the four biomarker regions reinforces these trends (**Fig. 2B**): in the sensitive strain, the 658∼663 cm¹ and 713∼719 cm¹ bands decrease progressively, whereas the 1430∼1450 cm¹ and 1589∼1595 cm¹ bands increase steadily (*p* < 0.0001). In resistant DH5α, early declines in the first two regions recover or stabilize, while the latter two exhibit brief reductions that subsequently plateau. Thus, both global spectra and biomarker dynamics enable robust discrimination between phage-sensitive and resistant phenotypes.

To assess detection sensitivity, we analyzed a weakly susceptible system: phage T4 infecting *E. coli* ATCC25922. Ramanomes were collected over 0∼100 min under MOI = 0, 1, and 10, with MOI at 0 as the control (**Fig. S1C, E**). In contrast, a strongly susceptible strain exhibits stable infection signatures even at MOI = 1 (**Fig. S2**), precluding limit-of-detection analysis; subsequent tests therefore focused on ATCC25922. At the global spectral level, infection-induced changes appear by 20 min at MOI = 10, but only after 80 min at MOI = 1. Analysis of the four biomarker regions (0, 40, 80, and 100 min) confirms this dependence (**Fig. 2C**). At MOI = 10, the 658∼663 and 713∼719 cm¹ bands decrease, whereas the 1430∼1450 and 1589∼1595 cm¹ bands increase consistently (*p* < 0.0001). At MOI = 1, these responses are delayed and temporally variable: the first two bands decline at 40 min (*p* < 0.05), rise at 80 min (*p* < 0.05), and sharply decrease again by 100 min (*p* < 0.0001); the latter two increase at 40∼80 min (*p* < 0.0001) but fall by 100 min (*p* < 0.0001). Thus, the diagnostic performance of individual Raman biomarkers depends on infection synchrony and population composition.

### Constructing a Composite Infection Index via Multi-Feature Integration for Rapid and Robust Infection Discrimination

To achieve robust RPST, the four Raman biomarkers were integrated into a unified diagnostic metric, called Composite Infection Index (CII), and tested using a weakly susceptible system, the T4-infecting *E. coli* ATCC25922. Spectra from uninfected and infected populations were labeled accordingly, and standardized intensities from the four biomarker regions (658∼663, 713∼719, 1430∼1450, and 1589∼1595 cm¹) were used as input features.

To identify the optimal integration strategy, we compared six approaches: direct summation of normalized biomarker intensities, logistic regression, linear discriminant analysis (LDA), naïve Bayes, support vector machine (SVM), and random forest (RF). Receiver operating characteristic (ROC) and precision-recall (PR) analyses showed that all machine-learning approaches achieve high performance (AUC > 0.99, accuracy > 94%), whereas direct summation performs poorly (AUC = 0.439 ± 0.040, accuracy = 0.454 ± 0.026; **Fig. 3A-B**). Among the classifiers, RF exhibited the highest ROC-AUC and PR-AUC values and was therefore selected for subsequent analyses.

**Figure 3.**
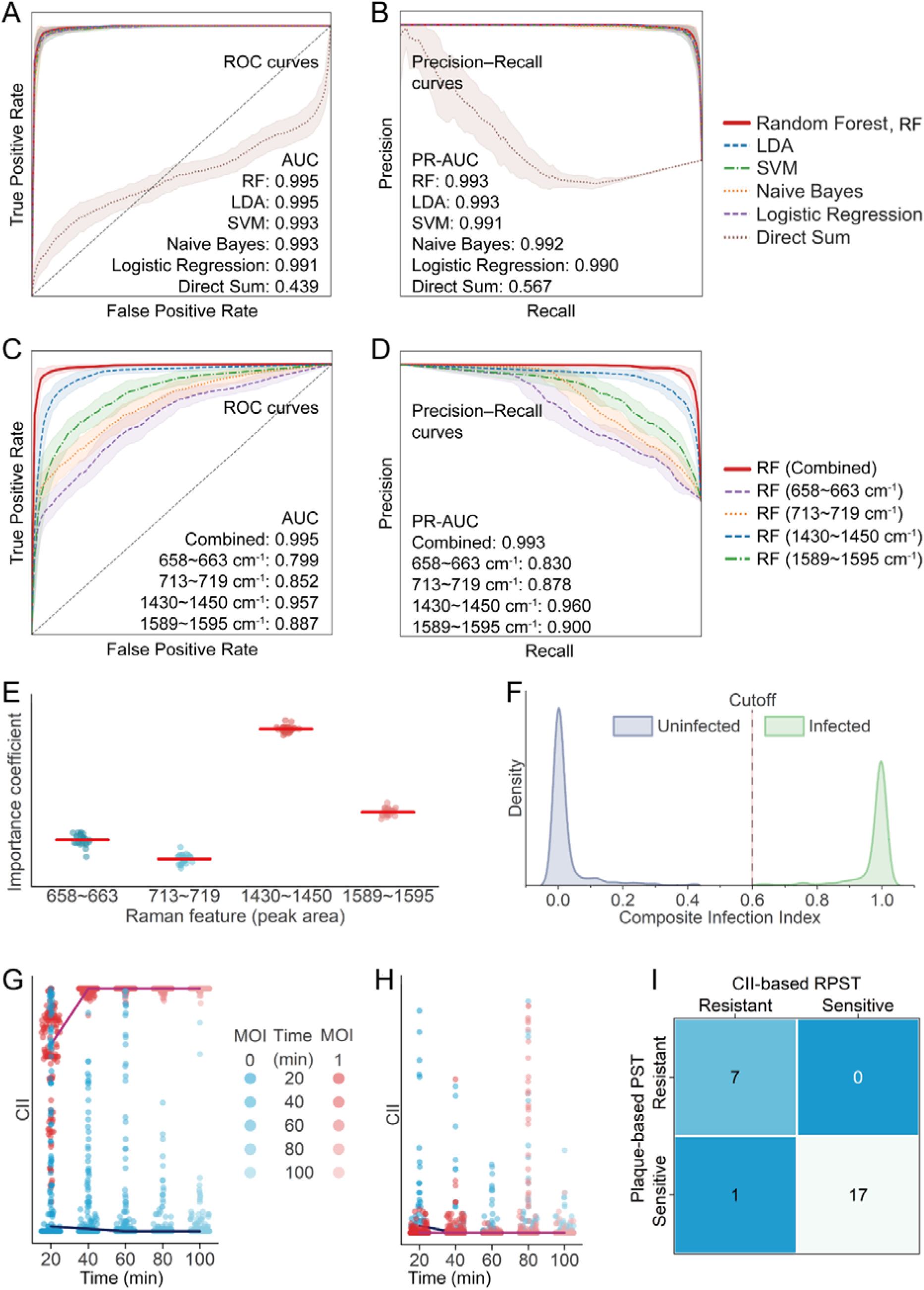
Construction of a composite infection index (CII) via multi-feature integration to achieve rapid and robust infection discrimination. (**A-B**) Performance comparison of six machine learning models integrating four RPST-derived biomarkers, evaluated by 5×5 cross-validation: direct summation, logistic regression, LDA, naïve Bayes, SVM, and RF. Based on (**A**) ROC and (**B**) precision-recall curves, both AUC and PR-AUC follow RF > LDA > SVM > naïve Bayes > logistic regression > direct summation, with RF achieving the best performance (AUC: 0.995 ± 0.003; PR-AUC: 0.993 ± 0.003). (**C-D**) Comparison between single biomarkers and the RF-based CII. Across repeated cross-validation, CII consistently outperforms all individual features in both (**C**) ROC and (**D**) precision–recall analyses. (**E**) RF-derived feature importance averaged across folds: 1430∼1450 cm¹ (0.516 ± 0.010), 1589∼1595 cm¹ (0.244 ± 0.012), 713∼719 cm¹ (0.151 ± 0.018), and 658∼663 cm¹ (0.089 ± 0.014). (**F**) Dual-cutoff framework for CII-based classification. The optimal cutoff (0.6000) maximizing G-mean defines infection status with 99% confidence bounds (lower: 0.5970; upper: 0.6030), categorizing populations as uninfected (CII < lower), infected (CII > upper), or uncertain (between cutoffs). (**G-H**) Temporal dynamics of CII. In T1-*E. coli* ATCC11303 (susceptible), CII exceeds the cutoff by 20 min (0.73 ± 0.21), stabilizing by 40 min (0.99 ± 0.02). In T1-*E. coli* DH5α (resistant), both the infected and control groups remain below the cutoff. (**I**) Validation across 25 phage-host systems (*E. coli*, *P. aeruginosa*, *Salmonella*, *K. pneumoniae*). CII-based classification agrees with plaque assays in 24/25 cases (96.0% accuracy).

To rigorously evaluate RF performance, we implemented a five-time repeated five-fold cross-validation scheme (25 total folds), yielding a mean AUC of 0.995 ± 0.003 and a mean accuracy of 0.965 ± 0.014. However, when trained on individual biomarkers, RF models show substantially lower performance (AUC = 0.80∼0.96; accuracy = 0.72∼0.89) compared with the integrated model (**Fig. 3C-D**), indicating that integration of multiple features improves classification robustness.

To clarify the basis of the performance gain, we examined the weight of RF features. All four biomarkers contribute to classification with distinct weights: 0.516 ± 0.010 (1430∼1450 cm¹), 0.244 ± 0.012 (1589∼1595 cm¹), 0.151 ± 0.018 (713∼719 cm¹), and 0.089 ± 0.014 (658∼663 cm¹) (**Fig. 3E**). The rankings remain consistent under various cross-validation folds (standard deviation of ranks < 0.2), confirming robustness of the selected biomarkers.

To implement CII as a diagnostic readout, ROC analysis was used to define a dual-cutoff framework for the CII. An optimal threshold (0.600) was identified by G-mean maximization and refined into low (0.597) and high (0.603) cutoffs at the 99% confidence level, yielding three outcome categories: uninfected (< low cutoff), infected (> high cutoff), and uncertain (between cutoffs) (**Fig. 3F**).

To validate diagnostic performance, we applied the CII-based PST to phage T1 infection of *E. coli* ATCC 11303 (susceptible) and DH5α (resistant) at MOI = 1. In ATCC11303, the CII rapidly increases, exceeding the infected cutoff by 20 min and stabilizing by 40 min (20 min: 0.73 ± 0.21; 40 min: 0.99 ± 0.02; 60∼100 min: ≥ 0.99 ± 0.01) (**Fig. 3G**). In contrast, DH5α consistently remains within the uninfected zone (**Fig. 3H**).

To evaluate generalizability, the CII was validated across 25 phage-host pairs encompassing *E. coli*, *Pseudomonas aeruginosa*, *Salmonella enterica*, and *Klebsiella pneumoniae*. After 40 minutes of co-incubation at MOI = 10, CII-based classification was compared with plaque assay results. The CII achieves concordant susceptibility calls in 24 of 25 systems (96.0% accuracy) (**Fig. 3I, Table S2**).

### Quantitation of Infection Efficacy for Rational Selection of Phage for Therapy

Beyond binary classification of PST, the continuous nature of the Composite Infection Index (CII) enables quantitative assessment of infection responses. Unlike plaque-based efficiency of plating (EOP), which yields discrete or qualitative outcomes, the CII provides a population-level metric that varies continuously with infection conditions.

To examine the dependence of CII on phage dosage, we analyzed the Ecp54-*E. coli* DH5α pair, across five MOIs (0, 0.01, 0.1, 1, and 10) after 40 min of co-incubation. CII values increase with MOI and exceed the infection cutoff (CII > 0.6030) at MOI ≥ 0.1 (**Fig. 4A**). Distribution analysis revealed that the fraction of infected cells rose progressively with MOI, exceeding uninfected cells at MOI = 0.1 and approaching complete infection at MOI = 10 (**Fig. 4B**). Thus, CII-based RPST can resolve continuous, MOI-dependent infection responses.

**Figure 4.**
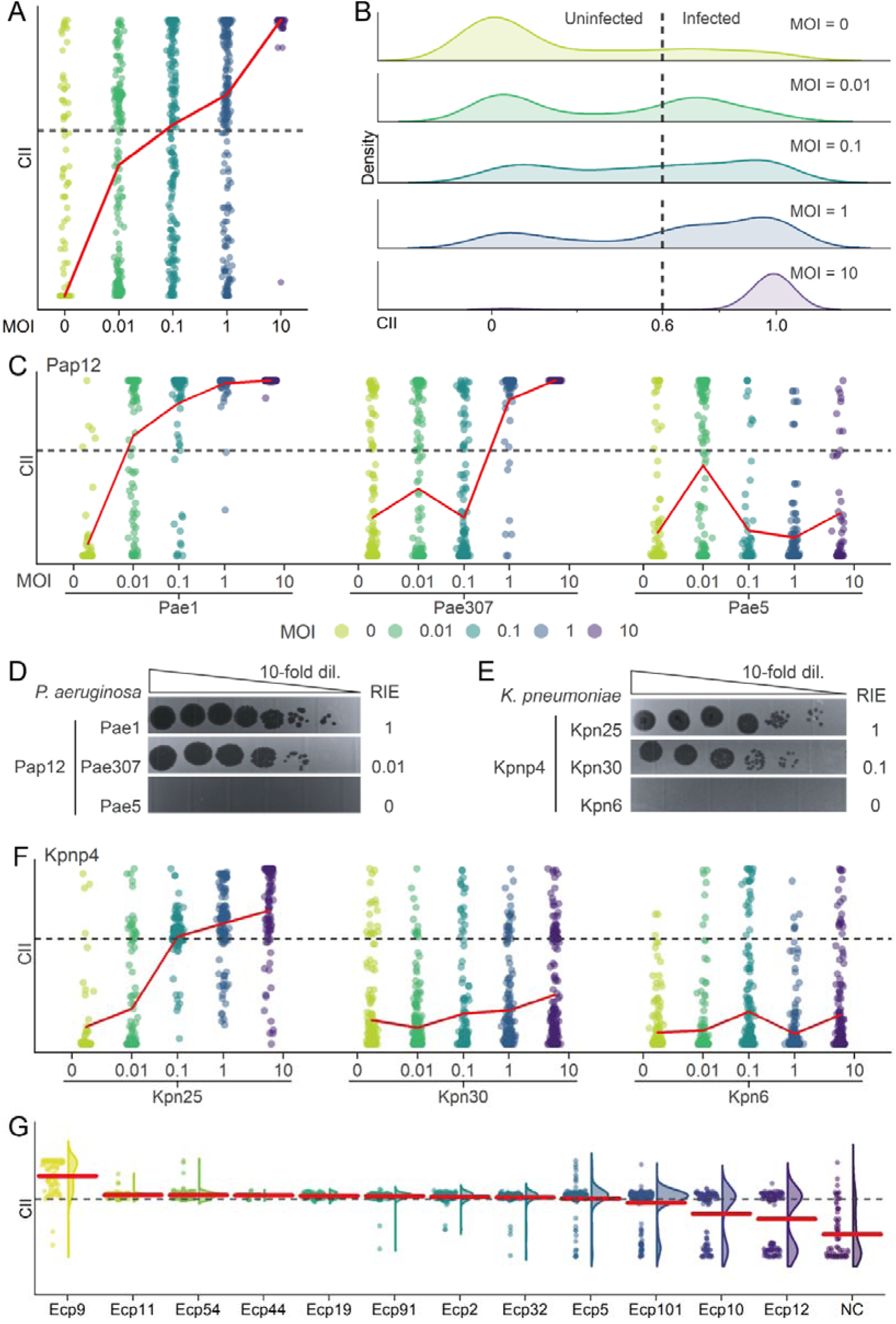
Quantitative assessment of relative phage potency by RPST. (**A-B**) Effect of multiplicity of infection (MOI) on the composite infection index (CII) in the Ecp54-*E. coli* DH5α system. (**A**) Boxplots show progressive CII elevation with increasing MOI after 40 min co-incubation. (**B**) Density plots at 40 min reveal a gradual rise in the proportion of infected cells, exceeding uninfected populations at MOI = 0.1 and approaching complete infection at MOI = 10. (**C-D**) Evaluation of phage Pap12 against three *P. aeruginosa* strains. (**C**) RPST-based CII distributions show MOI-dependent increases for susceptible strains Pae1 and Pae307, surpassing the infection cutoff at MOI = 0.01 and 1, respectively, while the resistant strain Pae5 remains below the cutoff even at MOI = 10. (**D**) Plaque-based relative infection efficiency (RIE) assays yield RIE = 1 (Pae1), 0.01 (Pae307), and 0 (Pae5). (**E-F**) Evaluation of phage Kpap4 against three *K. pneumoniae* strains. (**E**) RIE assays show values of 1, 0.1, and 0 for Kpn25, Kpn30, and Kpn6, respectively. (**F**) Corresponding RPST-based CII distributions demonstrate cutoff crossing at MOI = 0.1 for Kpn25, persistent sub-cutoff levels for Kpn6 even at MOI = 10, and gradual but incomplete increases for Kpn30 (RIE > 0). (**G**) Comparative analysis of twelve *E. coli* phages (MOI = 10, 40 min). Nine elevated population CII above the cutoff, with Ecp9 showing the highest CII. Density plots exhibit unimodal distributions for infective phages, while three noninfective ones display bimodal patterns with population CII remaining below the cutoff.

To determine whether RPST can resolve phage relative infectivity across multiple hosts, we tested phage Pap12 with *P. aeruginosa* isolates and phage Kpnp4 with *K. pneumoniae* isolates, each system including two plaque-positive and one plaque-negative strain. For *P. aeruginosa*, Pae1 enters the infectious state at MOI = 0.01, Pae307 at MOI = 1, while Pae5 remains below the threshold even at MOI = 10 (**Fig. 4C**), consistent with Relative Infection Efficiency (RIE) values of 1, 0.01, and 0, respectively (**Fig. 4D**). For *K. pneumoniae*, Kpn25 crosses the infection threshold at MOI = 0.1, Kpn6 remains uninfected at MOI = 10, and Kpn30 shows gradual CII increases without crossing the infection threshold (**Fig. 4F**), despite being plaque-positive (RIE = 0.1) (**Fig. 4E**).

To evaluate whether RPST can rank the relative efficacy of multiple phages against the same host, twelve *E. coli* lytic phages were tested against *E. coli* DH5α at MOI = 10 for 40 min. Nine systems exceed the infection threshold (**Fig. 4G**), with Ecp9 showing the highest CII (*p* < 0.0001). Notably, systems below the threshold exhibit bimodal CII distributions, reflecting mixed infected and uninfected subpopulations (**Fig. 4F**). Consistently, phages with CII above the threshold produce transparent plaques, whereas those below the threshold yield turbid or no plaques (**Fig. S3**).

### Determining the Minimal Effective MOI via Dynamics of Ramanomes during Infection

Beyond endpoint susceptibility classification, time-resolved ramanome profiling enables assessment of whether infection can be maintained over time under different phage-to-bacterium ratios. Here, we define the minimal effective MOI operationally as the lowest initial MOI at which the Composite Infection Index (CII) rises above the infection threshold and remains stably elevated over the observation period.

To assess whether RPST can resolve infection dynamics and determine the minimal effective MOI, we established a multidimensional experimental matrix incorporating five MOIs (0, 0.01, 0.1, 1, and 10) and six time points (0∼120 min at 20-min intervals) across three phage-bacterium systems: T1-*E. coli* ATCC 11303, T4-*E. coli* ATCC 11303, and T4-*E. coli* ATCC 25922. In the T1-ATCC11303 system, CII increases rapidly with MOI (**Fig. 5A**). At MOI = 10, the population surpasses the infection threshold within 20 min (CII = 0.97 ± 0.07 > cutoff). At lower MOI, CII rises more slowly but reaches comparable infected levels within 60 min (CII = 0.98 ± 0.03 ∼ 0.99 ± 0.05), indicating that infection can be sustained even under sparse initial phage input. Based on these dynamics, the minimal effective MOI for this system lies below 0.01.

**Figure 5.**
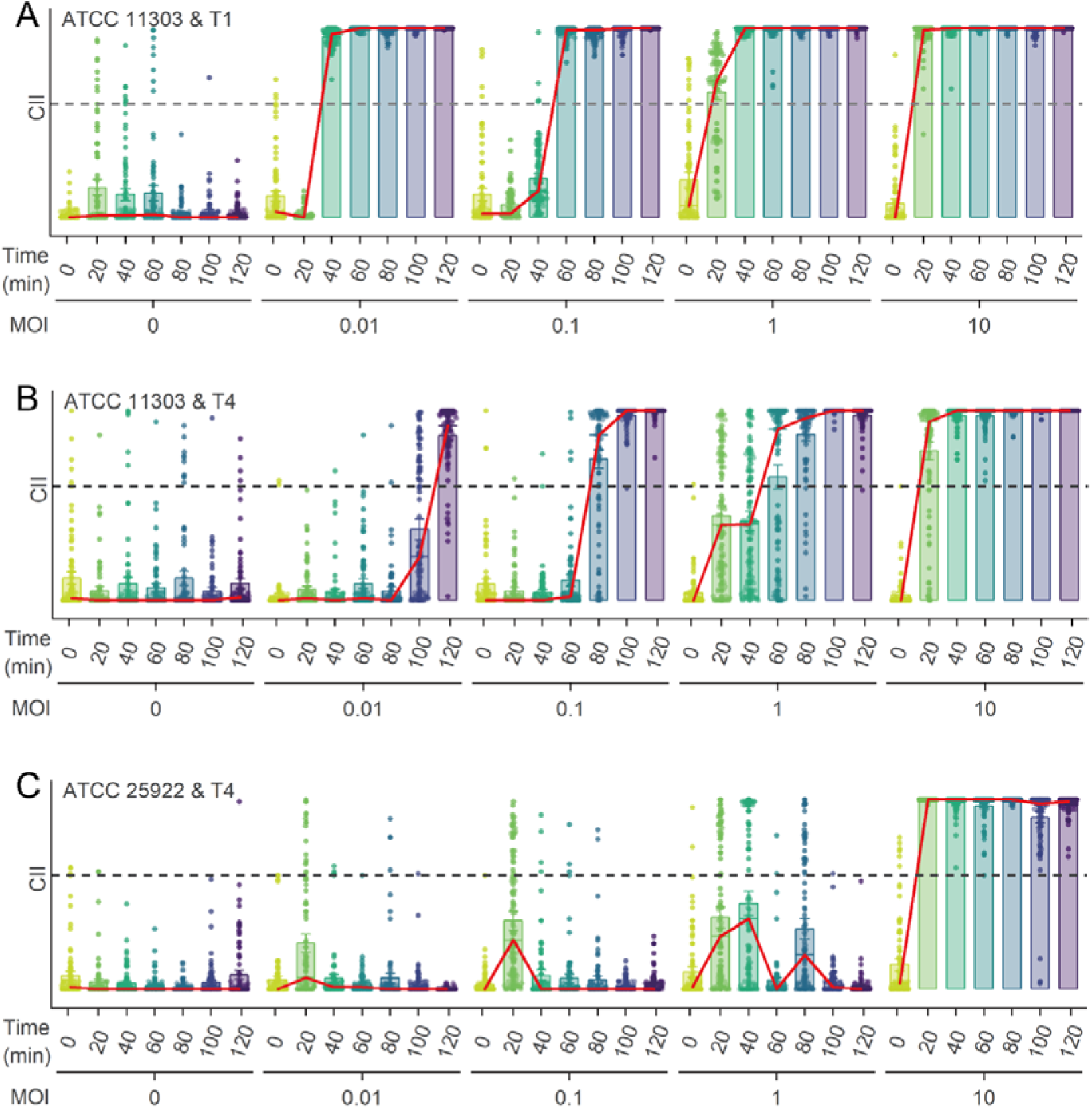
RPST enables quantitative assessment of effective MOI. (**A-C**) MOI- and time-resolved trajectories of CII in three phage-host systems. Populations were monitored under five MOIs (0, 0.01, 0.1, 1, and 10) across seven timepoints (0∼120 min, 20-min intervals). (**A**) In the T1-*E. coli* ATCC11303 system, populations at MOI = 0 remain below the infection cutoff throughout the experiment, whereas those at MOI = 10 cross the threshold within 20 min. Intermediate MOIs (0.01, 0.1, 1) exhibit delayed but complete transitions into the infected state within 60 min. (**B**) In the T4-*E. coli* ATCC11303 system, populations cross the infection cutoff within ≤ 20 min at MOI = 10, 60 min at MOI = 1, 80 min at MOI = 0.1, and 120 min at MOI = 0.01, while remaining uninfected at MOI = 0. (C) In the T4-*E. coli* ATCC25922 system, only MOI = 10 leads to a transition to the infected state. At lower MOIs, CII values rise transiently but subsequently decline, and population-level CII never exceeds the infection cutoff.

The T4-ATCC11303 system exhibits a similar qualitative pattern, with all MOI > 0 conditions eventually reaching stable infection (**Fig. 5B**). However, the transition kinetics differ markedly: higher MOI accelerates the onset of infection (≤ 20 min at MOI = 10), whereas lower MOI (0.01∼1) requires extended incubation (60∼120 min) before achieving comparable infection levels. Therefore, although both systems support infection propagation at low MOI, the temporal dynamics of establishment vary between phage-host pairs.

In contrast, in the T4-ATCC25922 system displays a distinct dynamic profile. Only MOI = 10 leads to sustained infection (CII ≈ 1 from 20∼120 min; **Fig. 5C**). At lower MOI (0.01∼1), CII exhibits transient increases followed by declines to baseline. For example, at MOI = 1, CII rises to 0.45 ± 0.41 by 40 min, falls to 0.04 ± 0.12 at 60 min, rises again to 0.32 ± 0.34 at 80 min, and returns to baseline thereafter, indicating that initial infection events have failed to propagate effectively within the population. Thus, initial infection events don’t lead to stable population-level infection, placing the minimal effective MOI for this system above 1. Together, time-resolved CII trajectories delineate system-specific thresholds for sustained infection and reveal pronounced differences in infection establishment kinetics across phage-host combinations.

## Discussion

Despite considerable progress in phage susceptibility testing (PST), current strategies remain unable to simultaneously meet the key diagnostic requirements of accurately determining host susceptibility, ranking lytic potency, and evaluating the effective MOI that can sustain infection. RPST addresses this gap by leveraging population-level Raman phenotyping as a direct, proliferation-independent readout of infection-induced biochemical remodeling.

At the population level, productive lytic phage infection entails a profound reallocation of host cellular resources, redirecting biosynthetic capacity from host maintenance toward viral genome replication and virion assembly (31–33). Consistent with the diagnostic objective of RPST, ramanome-based readouts capture relative, population-averaged biochemical remodeling rather than absolute molecular abundances. In fact, as Raman spectra were normalized by the total peak area across the fingerprint region, changes in individual bands reflect shifts in the relative composition of cellular constituents within infected populations. The four conserved Raman biomarkers map onto core biochemical remodeling processes, nucleic acid metabolism, protein biosynthesis, and CHC-rich macromolecular accumulation, that have been repeatedly identified as dominant host responses to lytic phage infection by independent omics studies across diverse systems (31–33). Specifically, the increase in the 1430∼1450 cm¹ band, often broadly assigned to lipids (34), more plausibly reflects enhanced protein-associated CHC vibrations arising from viral structural and replication-associated proteins rather than de novo lipid synthesis (35). Conversely, the decrease in the nucleic acid-associated 713∼719 cm¹ band does not indicate reduced nucleic acid synthesis, but rather that protein accumulation outpaces nucleic acid expansion during productive infection. Given the distinct Raman scattering cross-sections of different functional groups (36), individual spectral features are context-sensitive, underscoring the necessity of integrative, multi-biomarker strategies for robust and generalizable PST assessment.

However, infection-induced biochemical remodeling does not unfold synchronously across bacterial populations, particularly under low MOI or during early infection stages. Under these conditions, infected and uninfected cells coexist, and the fraction of productively infected cells evolves dynamically over time. As a result, even biologically conserved Raman features are vulnerable to dilution by uninfected or transiently perturbed subpopulations, leading to unstable or context-dependent diagnostic performance when considered in isolation.

Integrating multiple complementary Raman biomarkers into a composite infection index (CII) therefore provides a more stable and generalizable diagnostic signal. By aggregating biochemical dimensions associated with nucleic acids, proteins, and CHC-containing macromolecules, CII amplifies coherent infection-associated remodeling while suppressing noise arising from population heterogeneity. This integrative strategy enables robust discrimination across diverse phage-host systems, even under challenging conditions such as low MOI or asynchronous infection.

Crucially, CII-based RPST achieves high concordance with plaque-assay-based susceptibility calls across diverse phage-host systems and greatly reduces turnaround time. This establishes RPST as a reliable phenotypic PST readout capable of rapidly and accurately distinguishing susceptible from resistant bacterial strains at the population level.

Beyond binary susceptibility classification, the continuous nature of CII enables quantitative assessment of infection efficacy. Because CII reflects the fraction of cells undergoing sustained biochemical remodeling, its magnitude provides a population-level proxy for phage lytic potency under defined conditions. This property allows RPST to rank infective phages targeting the same host and to resolve differences that remain indistinguishable by plaque morphology alone. In contrast to plaque assays, which provide only qualitative or categorical readouts, RPST reframes phage susceptibility testing from a static compatibility assessment into a quantitative evaluation of infection strength, thereby supporting the rational selection of the most effective phage candidates.

Importantly, infection efficacy alone does not guarantee persistence. Effective phage therapy requires that infection dynamics be sustained under clinically realistic phage-to-bacterium ratios, such that phage amplification and secondary infection can outpace bacterial proliferation. By resolving CII trajectories across MOI and time, RPST quantifies the lowest phage-to-bacterium ratio that supports self-propagating infection at the population level, thereby defining the minimal effective MOI. Unlike plaque assays, which provide only endpoint compatibility, RPST captures both the initiation and persistence of infection, linking early phenotypic susceptibility with dynamic infection outcomes relevant to *in vivo* efficacy. Moreover, time-resolved RPST enables longitudinal monitoring of infection trajectories, revealing whether bacterial populations trend toward infection dominance or recovery. Although further validation is required, the minimal effective MOI derived from RPST provides a rational, quantitative metric to guide phage formulation and dosing under realistic infection conditions.

Nevertheless, there are several limitations. RPST prioritizes speed and throughput by profiling population-level Raman signals, which inevitably masks single-cell heterogeneity. Under low MOI or early infection stages, infected and uninfected cells may coexist, and infection-induced biochemical remodeling may fail to propagate across the population. In such cases, transient or intermediate CII values reflect non-sustained, population-level infection outcomes rather than definitive cellular fates. RPST therefore does not distinguish specific intracellular failure modes, such as abortive or pseudolysogenic infections (37), but instead reports whether infection-associated remodeling is self-sustaining at the population level. Future integration with single-cell Raman acquisition strategies, such as Raman Flow Cytometry (38), may enable more refined classification of infection states by resolving cell-to-cell heterogeneity that is inherently masked in population-level ramanome measurements. In addition, the generalizability of ramanome-based susceptibility metrics across instruments and laboratories remains an important consideration for clinical translation. RPST mitigates instrument-specific variability by relying on multiple conserved spectral bands spanning relatively broad wavenumber ranges, rather than single-wavenumber features, and by integrating these biochemical dimensions into the CII. Standard ramanome calibration procedures, including silicon-based wavenumber referencing and multi-peak molecular standards, can further support inter-platform reproducibility (39). While large-scale multi-center validation remains a future goal, these design choices provide a practical foundation for extending RPST across different Raman platforms. Moreover, although our validation has focused primarily on Gram-negative bacteria, the RPST biomarkers capture relative remodeling of core macromolecular categories rather than Gram-negative-specific structures. Consequently, the underlying diagnostic principle is, in principle, also applicable to Gram-positive organisms. On the other hand, differences in cell envelope architecture and baseline biochemical composition between Gram-negative and Gram-positive bacteria are expected to influence spectral baselines and infection dynamics. Accordingly, extension of RPST to Gram-positive species will require additional calibration and validation, rather than direct transfer of thresholds established here. Extending the spectral framework to Gram-positive organisms, as well as to polymicrobial samples and biofilm-associated infections, will therefore be an important step towards broader clinical translation. Besides, complex clinical matrices may introduce spectral interference and compositional variability, underscoring the need for systematic validation under increasingly realistic infection contexts.

With further optimization, RPST can substantially accelerate phage therapy development and clinical decision-making. Its rapid, label-free, and quantitative nature makes it suitable for not just preclinical phage screening, susceptibility testing, and potency ranking, but real-time monitoring of treatment response in personalized therapy and the rapid detection of emerging resistance. Looking forward, the integration of RPST with flow-mode (38, 40, 41) or multi-modal single-cell Raman-activated sorting (42, 43) and mass-parallelled, A.I.-directed single-cell culturomics (44, 45) can bridge the gap between phenotypic screening and the isolation of genetically unique phages or resistant bacterial clones, fueling discovery and mechanistic studies. More broadly, the RPST framework provides a generalizable paradigm for probing virus–microbe interactions through quantitative phenotypic fingerprints. By building expansive ramanomic databases, we can envision an AI-driven future where infection outcomes and optimal phage matches can be predicted directly from a bacterial isolate’s ramanome. Collectively, this work demonstrates that population-level Raman phenotyping can unify infectivity detection, potency quantification, and infection sustainability assessment within a single framework, paving the way to a more predictive and personalized approach to antimicrobial therapy.

## Materials and Methods

### Bacterial Strains, Media, and Culture Conditions

Reference strains *Escherichia coli* ATCC11303 and ATCC25922 were obtained from the American Type Culture Collection (ATCC, USA). The laboratory-stored *E. coli* DH5α and clinical isolates (*Salmonella enterica* Sal2, Sal8, Sal28; *Pseudomonas aeruginosa* Pae1, Pae5, Pae307; *Klebsiella pneumoniae* Kpn6, Kpn25, Kpn30) were provided by Shenzhen Third People’s Hospital (Shenzhen, China). All strains were cryopreserved at -80°C until use.

*E. coli* strains were cultured on Luria-Bertani (LB) agar and in LB broth (Hope Bio, China). *S. enterica* and *P. aeruginosa* strains were cultured on tryptic soy agar (TSA, Aobox, China) and in tryptic soy broth (TSB, Solarbio, China). *K. pneumoniae* strains were cultured on blood agar plates (Hope Bio, China) and in Mueller-Hinton broth (MHB, Hope Bio, China).

Before experiments, frozen strains were thawed, streaked onto respective agar plates, and incubated overnight at 37°C. Single colonies were inoculated into liquid media for subsequent phage-bacteria interaction experiments.

### Phage Types and Propagation

Lytic phages targeting *E. coli* (T1, T4, Ecp2, Ecp5, Ecp9, Ecp10, Ecp11, Ecp12, Ecp19, Ecp32, Ecp44, Ecp54, Ecp91, Ecp101), *S. enterica* (Salp6), *P. aeruginosa* (Pap12), and *K. pneumoniae* (Kpnp4) were obtained from Shenzhen Third People’s Hospital and stored at 4°C in SM buffer (50 mM Tris-HCl, 100 mM NaCl, 8 mM MgSOC, pH 7.5).

Phages were propagated on their respective laboratory hosts in liquid culture. Briefly, 100 µL of phage suspension (10^9^∼10^10^ PFU/mL) was added to 500 mL logarithmically growing bacteria (OD_600_ = 0.5) at a MOI of 0.01∼0.1. After incubation at 37 °C with shaking for 5 hours, lysates were centrifuged (10,000 × g for 30 min) and filtered (0.45 µm). Phage titers were determined by the double-layer agar method and expressed as plaque-forming units per milliliter (PFU/mL). Crude lysates were stored at 4 °C for short-term use or at −80 °C with 10% glycerol for long-term storage.

### Phage Sensitivity Assay (Double-Layer Agar Assay)

A 200 μL aliquot of mid-log-phase bacterial suspension (0.5 McFarland units) was mixed with 2 μL of phage suspension (10^8^ PFU/mL) in 5 mL of 0.7% semi-solid agar and overlaid onto respective agar plates. Plates were incubated at 37°C for 8 hours. All assays were performed in triplicate.

### Relative Infection Efficiency (RIE) Assay

Phage suspensions were serially diluted 10-fold in SM buffer, and 2 µL of each dilution was spotted on the bacterial lawn and cultured at 37°C overnight. The lowest phage titer producing visible plaques was recorded for each phage-host pair.

For each host, the phage requiring the lowest titer was defined as the reference (RIE = 1). The RIE of other phages was calculated as the ratio of the reference titer to the minimal plaque-forming titer of each test phage, reflecting their relative ability to initiate infection under identical conditions.

### Sample Preparation for Ramanome Detection

Bacterial colonies were suspended in respective liquid media and adjusted to 0.5 McFarland units (5 × 10^7^∼10^8^ CFU/mL) using a densitometer (530 nm). Phage suspensions were diluted to target MOIs and mixed with bacterial suspensions in equal volumes (n = 3 per group). Mixtures were incubated in 48-well plates at 37°C with shaking (200 rpm). At designated time points (0∼120 minutes, 20-minute intervals), 1 mL aliquots were centrifuged (10,000 × g, 2 minutes), washed thrice with ultrapure water, and resuspended in 10 μL sterile water. Approx. 3 μL of the suspension was placed onto quartz substrates for ramanome analysis.

### Raman Spectrometry

Spectra were collected using a confocal Raman microscope (RAMS, Qingdao Single-Cell Biotechnology Co., Ltd., China) equipped with a 532 nm laser, 1200 lines/mm grating, and a 100 × objective (NA = 1.25; Olympus, Japan). Prior to spectral acquisition, wavenumber calibration was performed using a silicon standard with a laser power of 20 mW, a pinhole size of 25 μm, and an integration time of 0.5 second. Acetaminophen was additionally measured as a molecular reference under a laser power of 20 mW, a pinhole size of 125 μm, and an integration time of 1 second.

Raman spectra of samples were then acquired with a laser power of 30 mW and an integration time of 1 second. Approx. 60 spectra were acquired per sample over the 400∼3500 cm¹ region, and the sampling depth is sufficient to accurately quantify metabolic features of the sample (28, 29).

### Ramanomic Data Preprocessing and Analysis

Raw spectra were performed using the RamEx package (version 1.0.0) (46), which integrates multiple processing modules, including cosmic-ray removal, spectral smoothing (Savitzky-Golay), baseline correction (polynomial fitting), spectral-range selection (fingerprint region), and peak area normalization. The *find_markers_roc* function (RamEx package: https://github.com/qibebt-bioinfo/RamEx) identified infection-specific biomarkers based on receiver operating characteristic (ROC) curves (AUC > 0.80). Statistical significance was determined using Student’s *t*-test, with *p* < 0.05 and 95% confidence intervals considered statistically significant unless otherwise specified.

### Random Forest Model Training and Composite Infection Index (CII) Calculation

A random forest (RF) classifier was implemented in Python (RRID:SCR_008394) 3.9 using the *scikit-learn* library (v 1.3.2). The model was trained on four Raman biomarkers from labeled infected and uninfected bacterial populations, with the predicted infection probability defined as the CII for each sample. The model utilized the following fixed hyperparameters to ensure reproducibility: n_estimators = 100, max_depth = None, min_samples_split = 2, min_samples_leaf = 1, criterion = ’gini’, bootstrap = True, and random_state = 42.

Model performance and generalizability were evaluated using a five-time repeated five-fold cross-validation scheme (25 total folds). In each iteration, four folds were used for training and one for testing, ensuring that every sample was included in a test fold. The classifier’s performance was assessed by AUC and accuracy computed across all test folds. Feature importance was computed for each fold to assess the stability of biomarker contributions.

ROC-based dual cutoffs were applied to classify samples as uninfected, infected, or uncertain. This framework enables reproducible CII computation for downstream analyses, including MOI-dependent infection dynamics and comparative phage potency.

### Additional Data Analysis and Visualization

Statistical analysis was performed using SPSS 18 (IBM, USA, RRID: SCR_002865). For group comparisons, Student’s *t*-test was employed with a nominal significance level of *p* < 0.05. All statistical tests were two-sided. Comparisons were defined a priori between specified experimental conditions at each time point, and different time points were treated as independent experimental conditions rather than repeated measures. Accordingly, no multiple comparisons were conducted on the same dataset, and no correction for multiple testing was applied. Data visualization was implemented with OriginPro 2021 (OriginLab Corporation, USA, RRID:SCR_014212), custom Python (RRID:SCR_008394) 3.09 scripts, and schematic illustrations created via BioRender.com.

## Data Availability

All data and code supporting the findings of this study are publicly available in the Zenodo repository. (*i*) The raw ramanome dataset is available at: https://doi.org/10.5281/zenodo.17470426; (*ii*) The Python source code for the Composite Infection Index (CII) calculation is available at: https://doi.org/10.5281/zenodo.17470774.

## Ethics statement

As all bacterial strains and phages were from residual samples used in clinical diagnosis or strains from their subcultures, the criteria for exemption were met.

## Acknowledgement

Special thanks are extended to Yang Liu and Zongyi Ma for their guidance on figure preparation.

## Author contribution

**Conceptualization**: J.X. and X.H.

**Supervision**: J.X. and X.H.

**Methodology**: X.H. (RPST development), X.W. (phage enrichment and co-incubation systems), X.F. and B.G. (Ramanome acquisition)

**Resources**: Y.Z. and H.L. (lytic phages and clinical bacterial isolates)

**Data Curation**: X.H.

**Formal Analysis**: X.H.

**Investigation**: X.W., X.F., B.G., Y.Z., and H.L.

**Writing-Original Draft**: X.H. and J.X.

**Writing-Review & Editing**: J.X., J.H., H.L., Y.Z., X.Z., X.H., S.H. and A.G.

**Project Administration**: J.X.

**Funding Acquisition**: J.X. and A.G.

All authors read and approved the final manuscript.

