## Supplementary Materials for "Rapid and Quantitative Phage Susceptibility Test by Ramanome"

**This PDF file includes:**

Supplementary Tables (S1, S2)

Supplementary Figures (S1 to S3)

References (1 to 10)

**Other Supplementary Materials for this manuscript include the following:**

Supplementary Data S1

### Supplemental Tables

**Supplemental Table 1. Biological assignments of SCRS.**

| Raman shift （$cm^{-1}$) | Biomolecule assignment | Molecular vibration | Reference |
|---|---|---|---|
| 666 | Protein | Cysteine C-S stretching vibration | (1) |
| 669 | Nucleic acids | Guanine ring breathing modes | (2) |
| 719 | Phospholipids | C-N stretching vibration | (3) |
| 723 | Nucleic acids | Adenine ring breathing mode | (4) |
| 780 | Nucleic acids | Ring breathing modes of cytosine, thymine, and uracil | (5) |
| 804 | Nucleic acids | Stretching vibrations of O-P-O | (6) |
| 843 | Glucose | CH rocking | (7) |
| 871 | Protein | Single bond stretching vibrations for the proline and valine | (8) |
| 993 | Protein | Phenylalanine ring breathing mode | (9) |
| 1023 | Glycogen | Carbohydrates Deformation vibration of C-O-H | (7) |
| 1094 | Nucleic acids | Symmetric phosphate stretching vibration | (5) |
| 1113 | Protein | Benzoid ring deformation | (8) |
| 1242 | Nucleic acids | C-N in-plane stretching | (3) |
| 1285 | Lipids | C=C stretching mode | (8) |
| 1330 | Nucleic acids | $CH_3CH_2$ wagging | (3) |
| 1369 | Phospholipids | $CH_3$ stretching mode | (4) |
| 1408 | Lipids | $CH_2$ stretching mode | (9) |
| 1444 | Lipids, protein | $CH_2$ scissoring mode | (6, 8) |
| 1471 | Lipids | $CH_2$ bending mode | (6) |
| 1476 | Nucleic acids | Nucleotide acid purine bases | (3) |
| 1525 | Carotenoid | C=C stretching mode | (10) |
| 1574 | Nucleic acids | Ring breathing modes of bases | (5) |
| 1595 | Protein | Phenylalanine C=C skeletal vibration | (6) |
| 1676 | Nucleic acids | Ring breathing modes of cytosine, thymine, and uracil | (8) |

**Supplemental Table 2. Comparison of S/R derived from Plaque assay and RPST.**

| Species | Strain | Phage | Sensitive (S) /Resistant (R) | | MEM_RPST* |
|---|---|---|---|---|---|
| | | | Plaque assay | RPST | |
| *E. coli* | ATCC11303 | T1 | S | S | 0.01 |
| | | T4 | S | S | 0.01 |
| | ATCC25922 | T4 | S | S | 10 |
| | DH5*α* | T1 | R | R | > 10 |
| | | Ecp2 | S | S | < 10 |
| | | Ecp5 | S | S | < 10 |
| | | Ecp10 | R | R | > 10 |
| | | Ecp11 | S | S | < 10 |
| | | Ecp19 | S | S | < 10 |
| | | Ecp44 | S | S | < 10 |
| | | Ecp91 | S | S | < 10 |
| | | Ecp101 | R | R | > 10 |
| | | Ecp9 | S | S | 0.1 |
| | | Ecp32 | S | S | 0.01 |
| | | Ecp54 | S | S | 0.1 |
| | | Ecp12 | R | R | > 10 |
| *S. enterica* | Sal8 | Salp6 | S | S | 1 |
| | Sal28 | | S | S | < 10 |
| | Sal2 | | R | R | > 10 |
| *P. aeruginosa* | Pae1 | Pap12 | S | S | 0.01 |
| | Pae307 | | S | S | 1 |
| | Pae5 | | R | R | > 10 |
| *K. pneumoniae* | Kpn25 | Kpnp4 | S | S | 0.1 |
| | Kpn30 | | S | R ϕ | > 10 |
| | Kpn6 | | R | R | > 10 |

* MEM_RPST: the minimum effective MOI of phages based on RPST.

ϕ Gray-shaded cells indicate cases where RPST and plaque assay results were discordant.

### Supplemental Figures

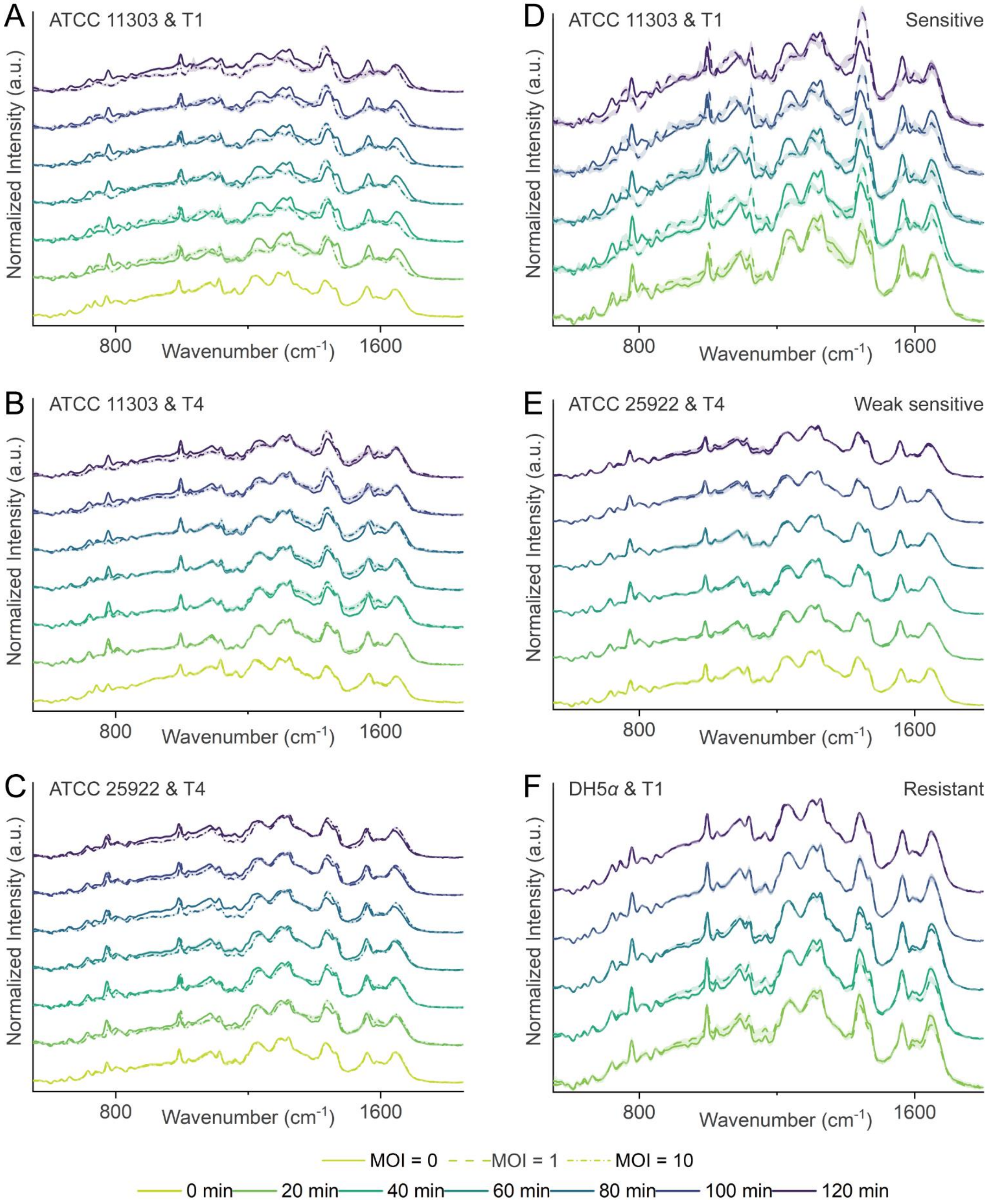


**Figure S1. Population-level Ramanome dynamics during phage–host co-incubation.** Ramanomes were collected from bacteria co-incubated with different phages to visualize overall spectral changes over time. (**A-C**) Spectral comparisons between infected (MOI = 10) and uninfected (MOI = 0) groups in three sensitive systems: (**A**) T1-*E. coli* ATCC11303, (**B**) T4-*E. coli* ATCC11303, and (**C**) T4-*E. coli* ATCC25922, measured at seven time points (0~120 min, 20-min intervals). (**D-F**) Spectral comparisons between experimental (MOI = 1) and control (MOI = 0) groups in three systems: (**D**) T1-*E. coli* ATCC11303, (**E**) T4-*E. coli* ATCC25922, and (**F**) T1-*E. coli* DH5$\alpha$, measured at five time points (20~100 min, 20-min intervals).

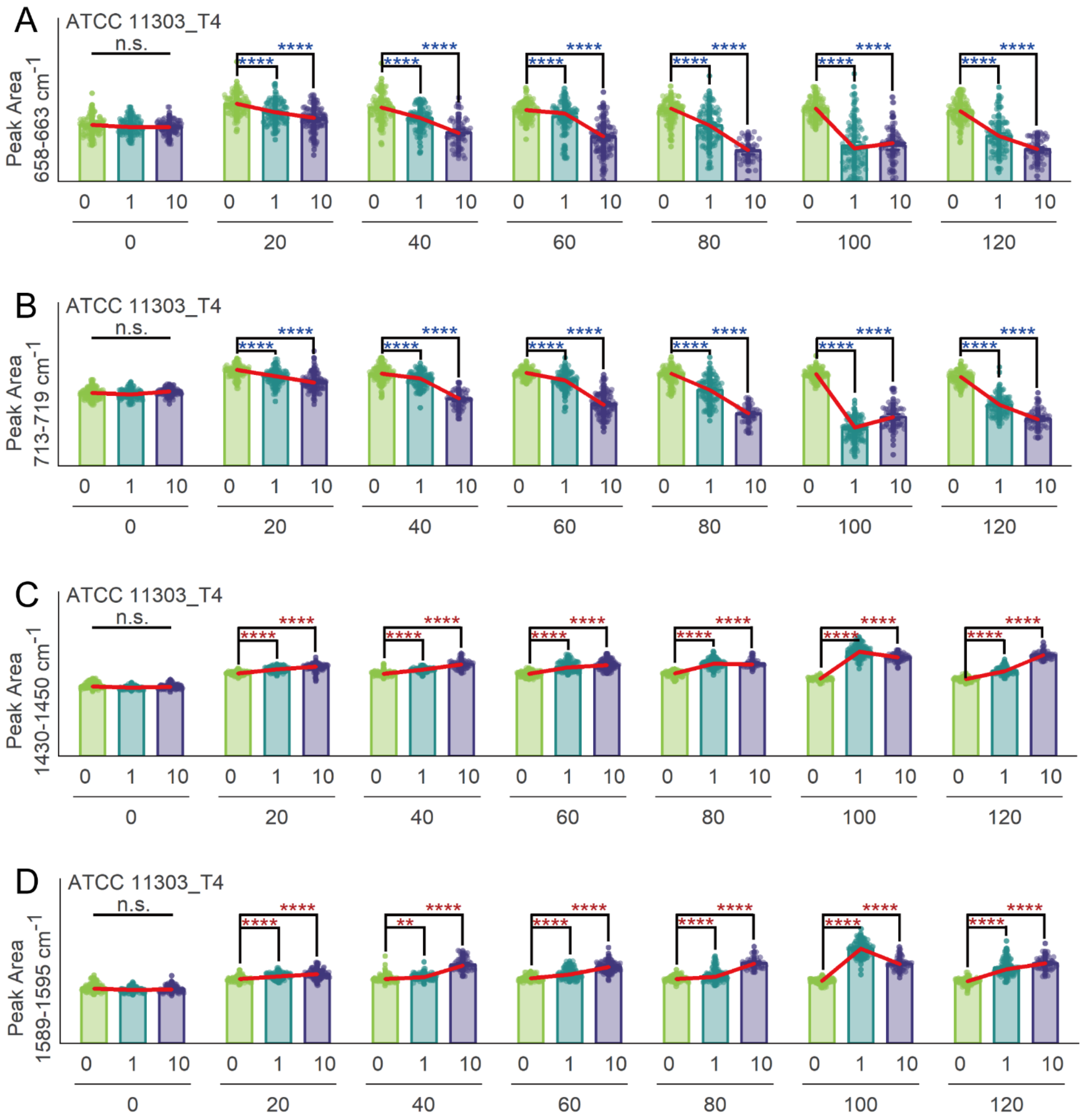


**Figure S2. Consistent time-dependent shifts of four Raman biomarkers in a highly sensitive system.** *E. coli* ATCC11303 were co-incubated with T4 phage at MOI = 0, 1, or 10, and ramanomes were collected at 0, 20, 40, 60, 80, 100, and 120 min. In the infected groups, the relative intensities of (**A**) 658~663 cm$^{-1}$ and (**B**) 713~719 cm$^{-1}$ decrease starting from 20 min post-infection, with the decline becoming more pronounced over time (observed at both MOI = 1 and MOI = 10). In contrast, the relative intensities of (**C**) 1430~1450 cm$^{-1}$ and (**D**) 1589~1595 cm$^{-1}$ begin to increase from 20 min post-infection, with the increase becoming more pronounced over time (also observed at MOI of 1 and 10). n.s., not significant; $*p < 0.05$, $**p < 0.01$, $***p < 0.001$, $****p < 0.0001$.

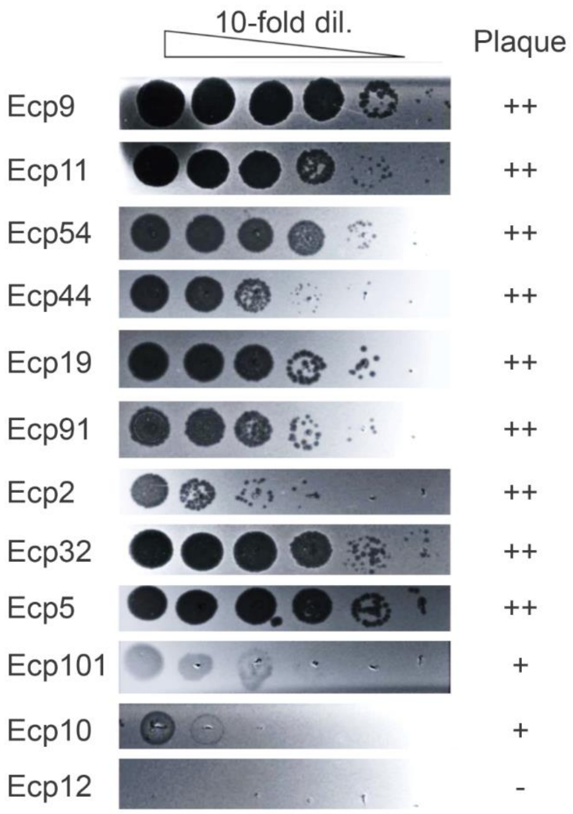


**Figure S3. Plaque assay outcomes of twelve lytic phages on *E. coli* DH5*α*.** Nine phages produce clear and well-defined plaques on *E. coli* DH5*α* via agar plates, whereas the other three yield turbid or indistinct plaques or no visible plaques at all.

**Data S1. (separate file)**

**Ramanome-based Composite Infection Index (CII) Dataset.**

This spreadsheet contains the analyzed ramanomic data for all bacterium-phage co-incubation systems investigated in this study. It reports the peak areas of the four characteristic Raman biomarkers used for the CII model, alongside the computed CII scores, predicted labels, confidence values, and predicted status for each spectrum based on the model output. The file also provides the final sample-level infection call, which is determined by aggregating the CII results from all ramanomes per sample.

The dataset is organized into multiple sheets: The first sheet, titled "Result_sum", provides a summary of the results at the sample level. Subsequent sheets are dedicated to the single-spectrum level data for each unique co-incubation system, named according to the convention: BacterialStrain_Phage (e.g., ATCC11303_T4).
